# Syndecan-4^-/-^ mice have smaller muscle fibers, increased Akt/mTOR/S6K1 and Notch/HES-1 pathways, and alterations in extracellular matrix components

**DOI:** 10.1101/2020.06.10.143982

**Authors:** Sissel Beate Rønning, Cathrine Rein Carlson, Jan Magnus Aronsen, Addolorata Pisconti, Vibeke Høst, Marianne Lunde, Kristian Hovde Liland, Ivar Sjaastad, Svein Olav Kolset, Geir Christensen, Mona Elisabeth Pedersen

## Abstract

**Background:** Extracellular matrix (ECM) remodeling is essential for skeletal muscle development and adaption in response to environmental cues such as exercise and injury. The cell surface proteoglycan syndecan-4 has been reported to be essential for muscle differentiation, but few molecular mechanisms are known. Syndecan-4^-/-^ mice are unable to regenerate damaged muscle, and display deficient satellite cell activation, proliferation, and differentiation. A reduced myofiber basal lamina has also been reported in syndecan-4^-/-^ muscle, indicating possible defects in ECM production. To get a better understanding of the underlying molecular mechanisms, we have here investigated the effects of syndecan-4 genetic ablation on molecules involved in ECM remodeling and muscle growth, both under steady state conditions and in response to exercise.

**Methods:** Tibialis anterior (TA) muscles from sedentary and exercised syndecan-4^-/-^ and WT mice were analyzed by immunohistochemistry, real-time PCR and western blotting.

**Results:** Compared to WT, we found that syndecan-4^-/-^ mice had reduced body weight, reduced muscle weight, muscle fibers with a smaller cross-sectional area, and reduced expression of myogenic regulatory transcription factors. Sedentary syndecan-4^-/-^ had also increased mRNA levels of syndecan-2, decorin, collagens, fibromodulin, biglycan, and LOX. Some of these latter ECM components were reduced at protein level, suggesting them to be more susceptible to degradation or less efficiently translated when syndecan-4 is absent. At the protein level, TRPC7 was reduced, whereas activation of the Akt/mTOR/S6K1 and Notch/HES-1 pathways were increased. Finally, although exercise induced upregulation of several of these components in WT, a further upregulation of these molecules was not observed in exercised syndecan-4^-/-^ mice.

**Conclusions:** Altogether our data suggest an important role of syndecan-4 in muscle development.

## Introduction

Skeletal muscle is a highly dynamic tissue, responding to physiological stimuli during development and exercise. It is composed of muscle cells (myofibers) collected in bundles. Both the individual myofibers and the myofiber bundles are surrounded by extracellular matrix (ECM). It was demonstrated more than 50 years ago that the satellite cells, the skeletal muscle stem cells (MuSCs), which are located between the basal lamina and sarcolemma (plasma membrane) of the skeletal muscle fibers and normally quiescent in adult muscle, become activated upon exercise, injury or disease, and are involved in the skeletal muscle regeneration (1). More recently, several reports have identified MuSCs as the primary contributors to the postnatal growth, maintenance and regeneration of skeletal muscles. These cells have a remarkable ability to self-renew, expand, or undergo myogenic differentiation to fuse and restore damaged muscle (2). The activation, proliferation and differentiation of MuSCs in adult muscle are mainly controlled by myogenic regulatory transcription factors such as myoblast determination protein (MyoD), myogenin, myogenic factor 5 (myf5) and 6 (myf6/mrf4). MuSCs express MyoD when they are activated and start to proliferate (3), whereas myogenin is crucial for later stages of myogenic differentiation (4).

The ECM, a complex network of collagens, in addition to glycoproteins and proteoglycans, is the niche of MuSCs and the myofibers. The small leucine rich proteoglycans (SLRPs) decorin, fibromodulin and biglycan, regulate collagen fibrillogenesis and crosslinking (5-7). The ECM composition is highly dynamic and promotes skeletal muscle adaption in response to environmental forces during e.g. exercise and regeneration, and during muscle growth and development. The ECM network also builds a scaffold for muscle cells to adhere, grow and differentiate, and functions as a storage and presenter of relevant muscle tissue growth factors and different cytokines during development (8).

Previous work has also identified syndecans to be important for muscle development, maintenance and regeneration (9-12). The syndecans belong to a group of transmembrane proteoglycans, which consists of four members in mammals (13). The syndecans are characterized by a large diverse extracellular domain with glycosaminoglycan (GAG) attachment sites, a conserved transmembrane domain and a short cytoplasmic domain with a unique variable domain differing between each syndecan. Syndecans have a wide spectrum of biological functions, and regulate calcium ion channels, polarization of epithelial cells, cell adhesions, and migration (14). They also function as co-receptors of various growth factors and transduce signals into the cell through their cytoplasmic domains (15). All four syndecans are expressed in developing muscles (16, 17) and in proliferating myoblasts, but their expression is progressively lost during myogenesis (18). Syndecan-2 is reported to be highly expressed in early-differentiated myoblasts (19). Syndecan-1 is not detected in postnatal muscle, while syndecan-3 and syndecan-4 are present and restricted to MuSCs and vascular cells (20, 21). Syndecan-4 as well as syndecan-3 have roles in development and regeneration. Both are highly expressed in myoblasts and around early embryonal myotubes but are reduced around myotubes postnatally and restricted to MuSCs in young adults (16). Syndecan-3 regulates muscle progenitor cell homeostasis by promoting MuSC self-renewal. Syndecan-3^-/-^ mice show improved muscle regeneration upon repeated muscle injuries, and reduced muscle pathology in dystrophic mice (11). Syndecan-4^-/-^ mice are unable to regenerate damaged muscle and display deficient MuSC activation, proliferation, myoD expression and differentiation (20). Interestingly, a reduced myofiber basal lamina has also been reported in syndecan-4^-/-^ muscle (20), indicating possible defects in ECM biosynthesis and turnover.

Limited knowledge on molecular mechanisms of syndecan-4 in skeletal muscle exist. To get a better understanding of the role of syndecan-4 in skeletal muscle and the underlying molecular mechanisms, we have here investigated the effects of syndecan-4 genetic ablation in tibialis anterior (TA), and analyzed molecules involved in ECM remodeling and muscle growth, both under steady state conditions and in response to exercise. Reduced muscle growth, changes in multiple signaling molecules and ECM components were found when syndecan-4 was absent. These molecular changes were mostly not affected further by exercise.

## Methods

### Animal experiments

All animal experiments were performed in accordance with the National Regulation on Animal Experimentation in accordance with an approved protocol (ID#2845 and 7696) and the Norwegian Animal Welfare Act and conform the NIH guidelines (2011). Female syndecan-4^-/-^ mice (KO) (22) were compared to either C57Bl/6j mice or syndecan-4^+/+^ bred from the same genetic background to study syndecan-4 dependent effects. Up to six mice per cage were housed in a temperature-regulated room with a 12:12-h light dark cycle, and access to food and water ad libitum. Animals were sacrificed by cardiac excision in deep surgical anesthesia.

### Exercise training protocol

Exercise training was performed on female syndecan-4^-/-^ mice (KO) and syndecan-4^+/+^ mice (WT) bred from the same genetic background with a treadmill for rodents (Columbus Instruments, OH, USA) with a 30-degree inclination. The mice were adapted two days to the treadmill, before exercise training was performed for 60 minutes per day for 14 days (one day rest without training) on a treadmill with moderate intensity. All mice had 3 minutes of warmup before each running session that consisted of 6 times 8 minutes of running followed by 2 minutes rest. Running speed was set to 14-18 meters per minute. Mice that were not able to complete the training protocol were excluded from the study. Animals were sacrificed by cardiac excision in deep surgical anesthesia.

### Real-time PCR

Tibialis anterior (TA) muscle was collected from WT and syndecan-4^-/-^ mice, exercised and sedentary mice, and snap-frozen in liquid nitrogen. Tissue samples were homogenized in lysis RLT-buffer of RNeasy minikit (#74104, Qiagen, Hilden, Germany) using a Precellys24 (#74106, Bertin Technologies, Villeurbanne, France) at 5500 rpm for 2×20 s, and RNA further purified following the manufacturer’s protocol of the RNeasy minikit including a DNase treatment on column according to the manufacturer’s protocol. cDNA was generated from ∼400 ng mRNA using TaqMan^®^ Reverse Transcription Reagents (Invitrogen, Carlsbad, CA, USA) according to the manufacturer’s protocol. The cDNA was diluted four times before aliquots (in duplicates) were subjected to real-time PCR analysis using an ABI Prism 7700 Sequence Detection system (Applied Biosystem, UK), and TaqMan^®^ primer/probe assays (see table 1 for primer/probes used) according to manufacturer’s protocol. The efficiency of each set of primers was always higher than 96%. Amplification of cDNA by 40 two-step cycles (15 sec at 95 °C for denaturation of DNA, 1 min at 60 °C for primer annealing and extension) was used, and cycle threshold (Ct) values were obtained graphically (Applied Biosystem, Sequence Detection System, Software version 2.2). ΔCt values and ΔΔCt values were calculated according to the MIQE guidelines (23). Comparison of the relative gene expression (fold change) was derived by using the comparative Ct method. In short, values were generated by subtracting ΔCt values between two samples which gives a ΔΔCt value. The relative gene expression (fold change) was then calculated by the formula 2^-ΔΔCt^ (24). Statistical analyses were performed using the ΔΔCt-values.

**Table 1.**
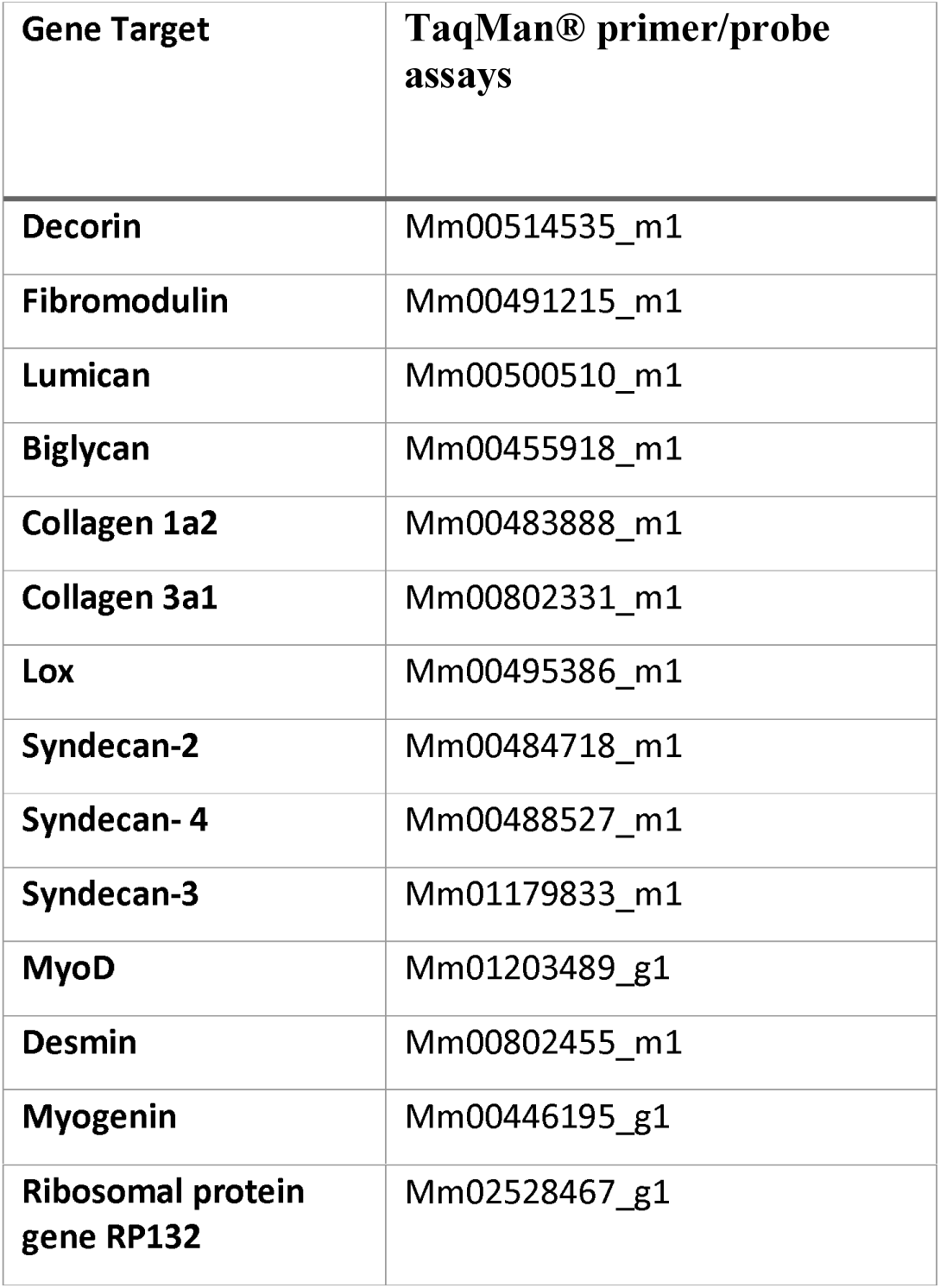
Gene target and TaqMan^®^ primer/probe assays.

### Immunofluorescence

TA muscles (n=4) were collected from WT and syndecan-4^-/-^ mice, embedded in OCT compound (Tissue Tek, Sakura Finetek, CA, USA) and then snap-frozen. Five micrometer-thick sections were cut on a cryostat and mounted on poly-L-lysine coated glass slides. The sections were air-dried for 5 min, before fixation using ice-cold acetone for five minutes. The sections were washed briefly twice in PBS, permeabilized using 0.5% Triton-X in PBS and incubated with 5% non-fat dry milk for 30 min before incubation with primary antibody for 1 h. Subsequent incubation with secondary antibodies was performed for 30 min before mounting using Dako fluorescent mounting medium (Glostrup, Denmark). Antibodies used were: Rabbit anti-Collagen 1 (ab34710, Abcam, Cambridge, UK), anti-pSer473-Akt (#9271, Cell Signaling, USA) and Alexa 488-goat anti-rabbit (#A-11078 Thermo Fisher Scientific, MA, USA). To quantify myofiber number sections were air-dried for 30 min and incubated with P-Block (5% goat serum, 1% Triton-X, 0.012% BSA, 0.012% non-fat dry-milk in PBS) for 1h. The sections were washed briefly four times with P-Block before incubation with primary antibody over night at 4 °C. Subsequent washing was performed twice with PBS before incubation with secondary antibody in P-Block for 1 h. The sections were washed briefly three times with P-Block, twice with PBS for five minutes each, rinsed in water before mounted using Dako fluorescent mounting medium. The total number of fibers in WT and KO TA muscles were quantified based on Laminin staining, using the ImageJ Cell counter plugin. Antibodies used were: Rabbit anti-Laminin (PA5-16287, Thermo Fisher Scientific, MA, USA). DyLight 549 mouse anti-rabbit were from Jackson Immunoresearch Cambridgeshire, UK. The sections were examined by fluorescence microscopy analysis (apotome mode) (ZEISS Axio Observer Z1 microscope, Jena, Germany), and images were processed using Adobe Photoshop CS3. If needed, brightness and contrast were adjusted manually across the entire image. Myofiber cross-sectional area in WT and KO TA muscles were quantified based on collagen staining. A minimum of 1,000 myofibers and four sections were analyzed per biological replica. Hoechst (Hoechst 33342, 10 µg/ml) was from Thermo Fisher Scientific. For assessment of connective tissue, sections were incubated with WGA Alexa Fluor™ 546 Conjugate (Thermo Fisher Scientific, MA, USA).

### Western blotting

Protein extracts from TA muscle were prepared using ice-cold lysis buffer (20 mm Hepes, pH 7.5, 150 mm NaCl, 1 mm EDTA, 0.5% Triton) supplemented with Complete EDTA-free protease inhibitor cocktail (#5056489001, Roche Applied Science, Merck, Darmstadt, Germany) and PhosSTOP (#4906837001, Roche Applied Science). Tissue samples were homogenized three times for 1 min on ice with a Polytron 1200 homogenizer and centrifuged at 14,000 × g for 10 min at 4 °C. Supernatants were collected and stored at −70 °C. Protein concentrations were determined using the Micro BCA protein assay kit (#23235, Pierce). The protein extracts were resolved on 4-15% Criterion™ Tris-HCl precast gels (#3450029, Bio-Rad Laboratories, CA, USA) and blotted onto PVDF membranes (#1704157, Trans-Blot Turbo Transfer Pack, Bio-Rad) using the Trans-Blot Turbo system (Bio-Rad) or tank blotting. The PVDF membranes were blocked in 5% non-fat dry milk, 1% casein or 3% BSA in TBS-T for 60 min at room temperature, followed by incubation with primary antibodies overnight at 4 °C. Membranes were washed two times for 10 min each in TBS-T, incubated with a horseradish-peroxidase-conjugated secondary antibody and thereafter washed two times for 10 min and one time for 5 min in TBS-T. Blots were developed using ECL Prime (RPN 2232, GE Healthcare, Il, USA). The chemiluminescence signals were detected by Las-4,000 (GE Healthcare). Membranes were re-probed after stripping using the Restore Western Blot Stripping buffer for 5 min at room temperature (21059, Thermo Fisher Scientific, MA, USA).

Antibodies and conditions were: anti-pSer235/236-RPS6 (1:1000, 5% BSA, #4858, Cell Signaling, MA, USA), anti-RPS6 (1:1000, 5% BSA, #2217, Cell Signaling), anti-HES-1 (1:500, 1% casein, #AB5702, Millipore, Merck, Darmstadt, Germany), anti-LRP6 (1:1000, 1% casein, #2560, Cell Signaling), anti-Dvl (1:1000, 1% casein, #3224, Cell Signaling), anti-β-catenin (1:5000, 1% casein, ab32572, Abcam, Cambridge, UK), anti-Frizzled-7 (1:1000, 1% casein, #ab64636, Abcam), anti-Cleaved Notch1 (1:1000, 5% BSA, #4147, Cell Signaling), anti-Wnt4 (1:1000, 1% casein, ab91226, Abcam), anti-TRPC7 (1:500, 1% casein, #SAB5200051, Sigma-Aldrich), anti-PKC (1:250, 1% casein, sc-208, Santa Cruz, Texas, USA), anti-DSCR1/RCAN1.4 (1:500, 5% milk, #D6694, Sigma-Aldrich), anti-pSer240/244-RPS6 (1:1000, 5% BSA, #5364 Cell Signaling), anti-pSer473-Akt (1:500, 5% BSA, #9271, Cell Signaling), anti-pThr308-Akt (1:500, #5106, 5% BSA, Cell Signaling), anti-Akt (1:500, 1% casein, #9272, Cell Signaling), anti-fibromodulin (1:500, 1% casein, sc-33772, Santa Cruz), anti-pSer2448-mTOR (1:1000, #2971, 5% BSA, Cell Signaling), anti-mTOR (1:1000, #2983, 5% BSA, Cell Signaling), anti-Pax-7 (1:500, sc-81648, 1:500, 1x casein, Santa Cruz) anti-decorin (1:500, 1% casein, AF1060, R&D Systems, MN, USA), anti-biglycan (1:500, 1% casein, ab49701, Abcam) and anti-LOX (1:500, 1% casein, sc-32409, Santa Cruz). Anti-rabbit IgG HRP (NA934V) affinity-purified polyclonal antibody, anti-mouse IgG HRP (NA931V) (both from GE Healthcare) and anti-goat IgG HRP (HAF109, R&D Systems MI, USA) were used as secondary antibodies. ProBlueSafeStain (Coomassie) was used to stain the membranes (#G00PB001, Giotto Biotech, FI, Italy).

### Statistics

All data were expressed as mean ±SEM. Comparisons between two groups were analyzed using Mann-Whitney U test (GraphPad Prism version 8.0.1, La Jolla, CA). A p-value of <0.05 was considered statistically significant. Comparison of fiber area sizes was performed using analysis of variance where each fiber’s estimated area was used as a sample, WT/KO was used as a fixed effect (the effect of interest) and magnification and muscle image number were included as random effects. The reason for including magnification was that image generation and fiber identification could be affected by the magnification. Muscle image was included to separate individual mouse effects from the true fiber area effects. Analyses were conducted in MATLAB 2015a, The MathWorks, Inc., Natick, Massachusetts, United States.

## Results

### Syndecan-4^-/-^ mice have reduced body weight, smaller muscle fibers and reduced myoD and myogenin mRNA levels

To investigate the skeletal muscle of syndecan-4^-/-^ mice, we analyzed the tibialis anterior (TA) muscles from adult female mice of 12-22 weeks. The syndecan-4^-/-^ mice were viable and showed no obvious postnatal abnormalities, however, they had a significantly lower body weight and TA weight compared to WT (Fig. 1A and 1B, respectively). Closer inspection showed that the syndecan-4^-/-^ muscle fibers had smaller cross-sectional area compared with those of the WT mice (Fig. 1C). More specifically, a larger fraction of the syndecan-4^-/-^ muscle fibers scored less than 2000 µm^2^ and there were hardly any larger muscle fibers (6000 µm^2^ and over) (Fig. 1D). Statistical analyses showed that the average size of the syndecan-4^-/-^ muscle fibers was 468 µm^2^ smaller compared to WT (p=0.0009), which was not accompanied by a change in total number of myofiber per muscle section (Fig.1E). As previously reported (20), the mRNA levels of the muscle transcriptional activator myoblast determination protein (myoD) and myogenin (this study) were significantly lower in syndecan-4^-/-^ compared to WT (Fig. 1F). Consistent with the previous report (20), we did not observe any centrally-nucleated myofibers in the syndecan-4^-/-^ (Suppl. Fig. 1A), a characteristic often associated with muscle disorders (20, 25) and impaired maintenance of MuSC quiescence (11). We did neither observe any differences in the protein levels of the paired box transcription factor Pax7, which is a marker of quiescent MuSCs (26) (Suppl. Fig. 1B). This is in line with Cornelison *et al*, who demonstrated no difference in the number of MuSCs in syndecan-4^-/-^ and WT (20).

**Fig. 1.**
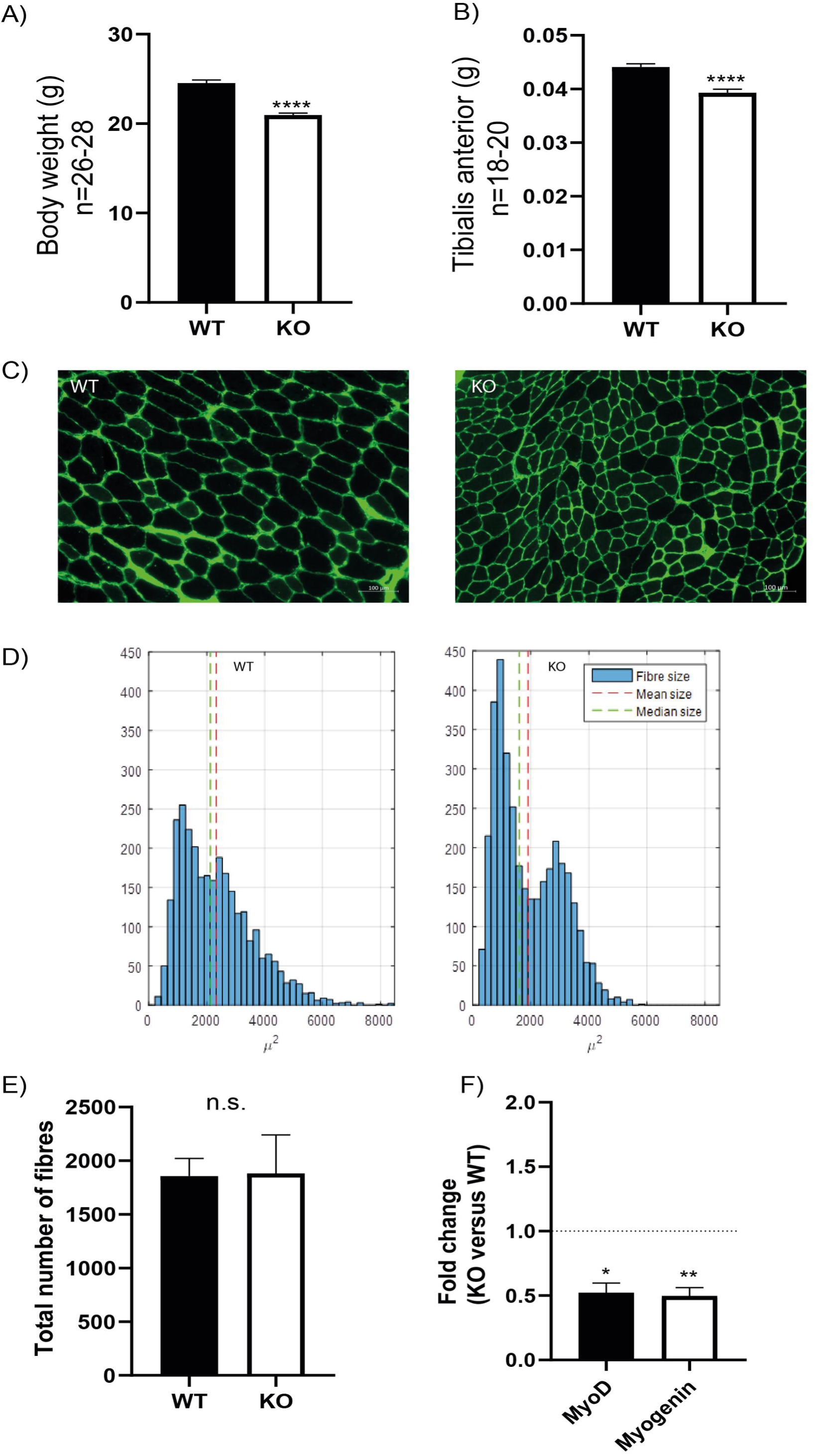
Syndecan-4^-/-^ mice have reduced body weight, smaller muscle fibers and reduced myoD and myogenin mRNA levels. Bars show (**A**) bodyweight and (**B**) tibialis anterior (TA) wet weight in syndecan-4^-/-^ mice compared to WT. The data are presented as average +/- SEM. Asterisks denote significant differences (***p<0.001) between syndecan-4^-/-^ and WT using un-paired two-tailed t-test. (**C**) Cross-sections of TA muscles from WT (left panel) and syndecan-4^-/-^ mice (right panel) were cryo-sectioned and stained with rabbit-anti collagen 1 (green), followed by Alexa Fluor 488-conjugated goat anti-rabbit before fluorescence microscopy analysis. (**D**) Quantification of images in **C** show fiber size distribution with a shift towards more fibers of low cross-sectional area (right panel) in syndecan-4^-/-^ compared to WT (left panel). Differences were tested using an analysis of variance with random effects. (**E**) Quantification of number of fibers in WT and syndecan-4^-/-^. The data are presented as average +/- SEM. Differences were tested using un-paired two-tailed t-test (ns; non-significant). **(F)** Bars show relative gene expression levels (fold change) in syndecan-4^-/-^ versus WT (young mice, aged 12-22 weeks, n=12) +/- SEM. The values of the bars are presented as fold change (in ΔΔCT) of syndecan-4^-/-^ related to WT (the latter is set to 1 and is represented as dotted line). Asterisk indicate significant differences, statistics assessed for ΔΔCT values by unpaired two-tailed t-test (*p<0.05, **p<0.01).

### The syndecan-4^-/-^ muscle has several changes in the ECM components

To further investigate the role of syndecan-4, syndecan-4^-/-^ and age-matched WT littermates were subjected to a two weeks-long treadmill exercise protocol (ET mice) (10-12 weeks at harvest). Sedentary syndecan-4^-/-^ and WT littermates of same age were used as controls (SED mice). As shown in Fig. 2A, the syndecan-4^-/-^ TA muscles were still significantly smaller than those from WT littermates after exercise (WT-ET versus KO-ET).

**Fig. 2.**
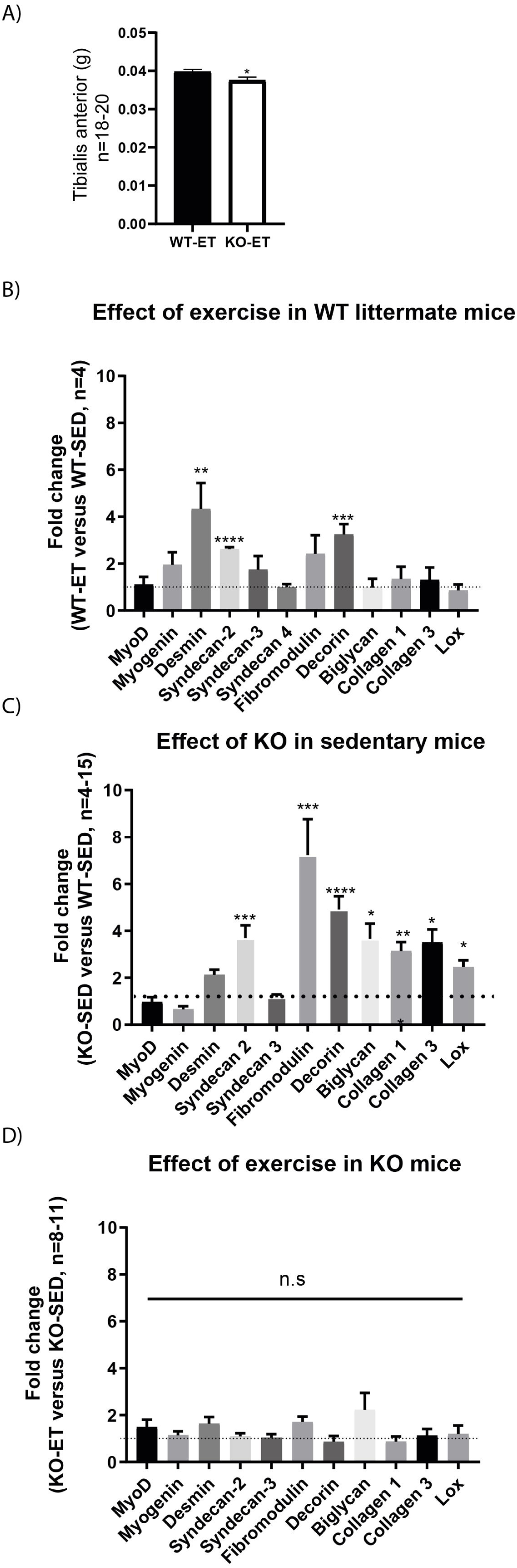
The syndecan-4^-/-^ muscle has increased mRNA levels of several ECM components. **(A)** Bars show tibialis anterior (TA) wet weight from exercised WT littermates (WT-ET) and syndecan-4^-/-^ mice (KO-ET) (10-12 weeks at harvest). The data is presented as the average +/- SEM. Comparisons between the groups were analyzed using Mann-Whitney U test (*p<0.05). Bars show relative mRNA levels in TA muscle from (**B**) WT-ET versus WT-SED, (**C**) KO-SED versus WT-SED and (**D**) KO-ET versus KO-SED (10-12 weeks at harvest). The values of the bars are presented as fold change (ΔΔCT) in WT-ET/WT-SED, KO-SED/WT-SED and KO-ET/KO-SED in **B, C** and **D**, respectively. WT-SED is set to 1 in **B** and **C**, whereas KO-SED is set to 1 in **D** (represented as dotted lines). Statistics are based on ΔΔCT values using unpaired two-tailed t-test (*p<0.05, **p<0.01, ***p<0.001).

To detect possible differences between syndecan-4^-/-^ and WT at the molecular level, we measured mRNA and protein levels of different ECM and signaling molecules we hypothesized could have a role in exercise and syndecan-4-mediated signaling. The mRNA levels of desmin and the ECM component decorin were both upregulated in WT after exercise (Fig. 2B, WT-ET versus WT-SED), which is consistent with a role of decorin and desmin in myoblast differentiation and fusion (18, 27). Syndecan-2 was also highly up-regulated in response to exercise, while no changes were seen in syndecan-3 and 4 (Fig. 2B). Myogenin and fibromodulin showed a tendency to increase in WT after exercise, whereas myoD, biglycan, collagen 1, collagen 3 and the collagen cross-linking enzyme lysyl oxidase (LOX) were unchanged (Fig. 2B). Consistently, although not significant, immunoblotting indicated a slight increased level of decorin and unchanged levels of fibromodulin, biglycan and LOX after exercise (Fig. 3A-D, WT-SED versus WT-ET).

**Fig. 3.**
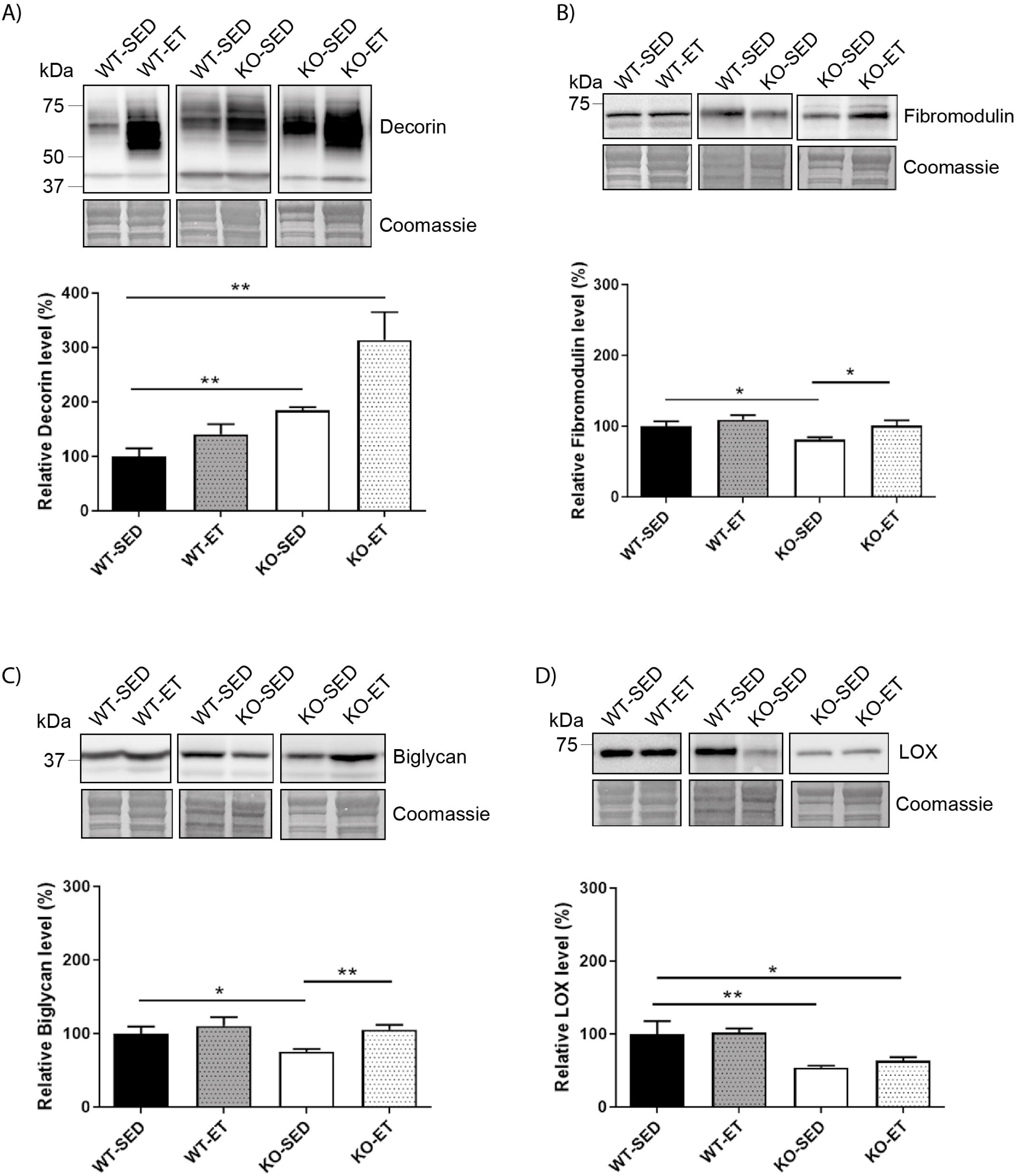
The syndecan-4^-/-^ muscle has changed protein levels of several ECM components. Immunoblot analyses of (**A**) decorin, (**B**) fibromodulin, (**C**) biglycan and (**D**) LOX in WT-SED, WT-ET, KO-SED and KO-ET. The values are presented in percentage and are normalized to WT-SED (n=4-10). Comparisons between the groups were analyzed using Mann-Whitney U test (*p<0.05, **p<0.01). ProBlue Safe Stain (Coomassie) was used as loading control.

Similar to exercise, syndecan-4^-/-^ loss led to increased mRNA levels of both syndecan-2 and decorin (Fig. 2C, KO-SED versus WT-SED). However, in contrast to WT, exercise failed to induce any further upregulation (Fig. 2D, KO-ET versus KO-SED). Interestingly, syndecan-4 loss also led to increased mRNA levels of ECM components and modifiers such as fibromodulin, biglycan, collagen 1, collagen 3 and LOX (Fig. 2C, KO-SED versus WT-SED). Consistently with the mRNA data, immunoblotting showed increased levels of decorin when syndcan-4 was absent (Fig. 3A, KO-SED versus WT-SED), but surprisingly a reduction in the fibromodulin, biglycan and LOX protein levels (Fig. 3B-D, KO-SED versus WT-SED). Notably, the biglycan and LOX protein levels increased in syndecan-4^-/-^ mice in response to exercise (Fig. 3C and D, KO-ET versus KO-SED), where fibromodulin and biglycan returned to the level observed in WT sedentary mice (Fig. 3B-C, KO-ET versus WT-SED).

Taken together, the syndecan-4^-/-^ muscle had increased mRNA levels of syndecan-2, collagen 1 and 3, decorin, fibromodulin, biglycan, and LOX. These three latter ECM components were reduced at protein level, suggesting that they might be more susceptible to degradation or less efficiently translated when syndecan-4 is absent. Although exercise induced upregulation of desmin, syndecan-2 and decorin in WT mice, a further upregulation was not observed in exercised syndecan-4^-/-^ mice.

### The syndecan-4^-/-^ muscle has increased Akt/mTOR/S6K1 and Notch-HES-1 pathways

Next, we analyzed activation of the protein kinase Akt, which transduces decorin-insulin-like growth factor I receptor (IGF1R) signaling (28) and regulates muscle fiber growth and hypertrophy through the Akt/mTOR/S6K1 pathway (29). Immunofluorescence data showed that pSer473-Akt localized to the skeletal muscle cell nuclei and to the ECM perimysium in WT where fibroblasts and resident immune cells are located (Fig. 4A, upper panels). Visual inspection clearly showed an increased expression of pSer473 in syndecan-4^-/-^ (Fig. 4A, lower panels). Consistently, immunoblotting revealed that the pSer473-Akt levels were constitutively higher in both sedentary as well as exercised syndecan-4^-/-^ mice compared to WT mice (Fig. 4B, KO-SED and KO-ET versus WT-SED). Also the levels of pThr308-Akt/Akt and pSer2448-mTOR/mTOR showed a tendency to be increased in sedentary syndecan-4^-/-^ mice (Suppl. Fig. 2A and Fig. 4C, KO-SED versus WT-SED). We also analyzed the phosphorylation levels of the 40S ribosomal protein S6 (RPS6), which is downstream of the Akt/mTOR/S6K1 signaling, and directly links to cell size control and myofiber growth (30, 31). Immunoblotting revealed that the pSer235/236-RPS6 levels were increased in WT muscles upon exercise (Fig. 4D, WT-ET versus WT-SED), but also in sedentary and exercised syndecan-4^-/-^ (Fig. 4D, KO-SED and KO-ET versus WT-SED). The levels of pSer240/244-RPS6 were also slightly increased in syndecan-4^-/-^ muscles, although the increase was not statistically significant (Suppl. Fig. 2B, KO-SED versus WT-SED). No difference in the total RPS6 levels was observed between syndecan-4^-/-^ and WT muscles (Fig 4D, WT-SED versus KO-SED, lower most panel).

**Fig. 4.**
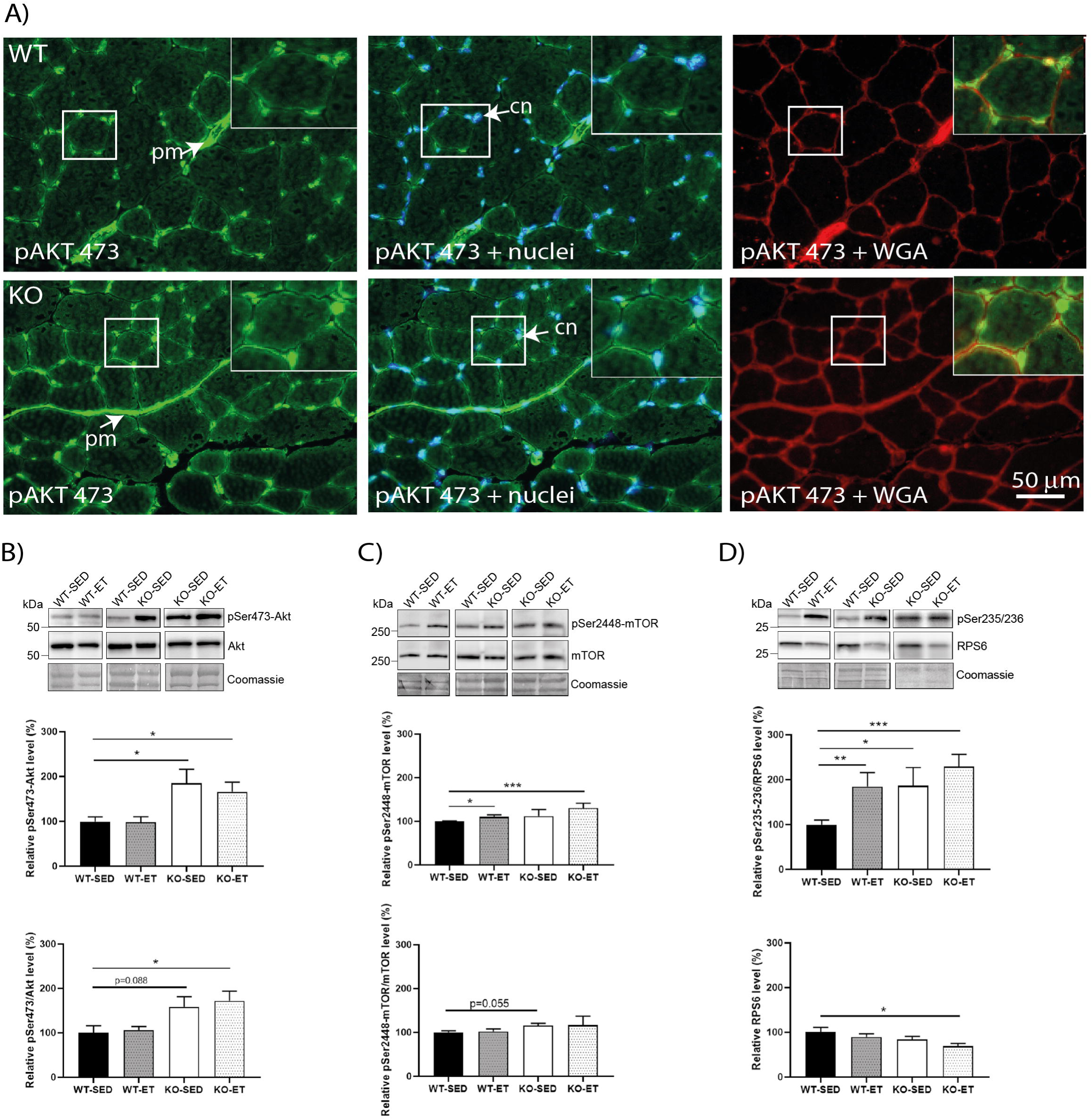
The syndecan-4^-/-^ muscle has an increased Akt/mTOR/S6K1 pathway. **(A)** Immunofluorescence of pSer473-Akt (green) in WT and syndecan-4^-/-^ muscles. The cell nuclei were counterstained with Hoechst (blue). The connective tissue was stained with WGA (red). *cn* cell nuclei, *pm* perimysium, indicated with arrows. Inserts show magnification of boxed areas. Scalebar as indicated. Immunoblot analyses of (**B**) pSer473-Akt (upper panels) and Akt (middle panels), (**C**) pSer2448-mTOR (upper panels) and mTOR (middle panels), and (**D**) pSer235/236-RPS6 (upper panels) and RPS6 (middle panels) in WT-SED, WT-ET, KO-SED and KO-ET. The values are presented in percentage and are normalized to WT-SED (n=6-10). Comparisons between the groups were analyzed using Mann-Whitney U test (*p<0.05, **p<0.01). ProBlue Safe Stain (Coomassie) was used as loading control (lower panels in B-D).

We next analyzed Cleaved Notch, as Notch signaling associates with increased syndecan-2 levels (32), and is known to increase upon exercise and be involved in myogenesis (33). Although our data indicated no significant changes of Cleaved Notch1 in WT (Fig. 5A, WT-ET versus WT-SED), the Notch target gene HES-1 was increased in response to exercise (Fig. 5B, WT-ET versus WT-SED). Surprisingly, both the Cleaved Notch and HES-1 levels were increased in sedentary syndecan-4^-/-^ mice and did not further increase in response to exercise (Fig. 5A-B, KO-SED versus WT-SED).

**Fig. 5.**
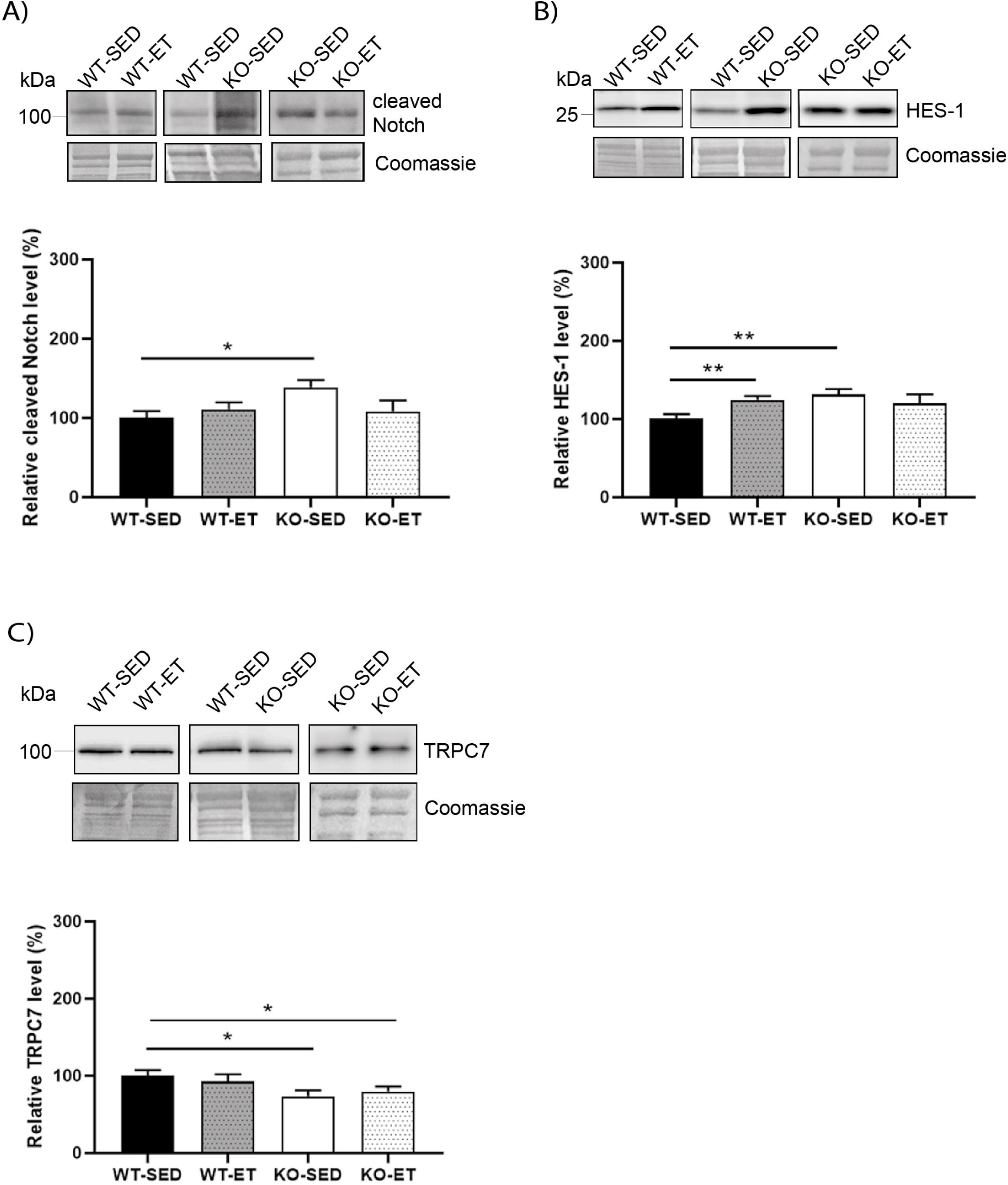
The syndecan-4^-/-^ muscle has an increased Notch/HES-1 pathway and a reduced TRPC7 level. Immunoblot analyses of (**A**) Cleaved Notch, (**B**) HES-1, and (**C**) TRPC7 in WT-SED, WT-ET, KO-SED and KO-ET. The values are presented in percentage and are normalized to WT-SED (n=6-10). Comparisons between the groups were analyzed using Mann-Whitney U test (*p<0.05, **p<0.01). ProBlue Safe Stain (Coomassie) was used as loading control (lower panels in A-C).

Finally, since syndecan-4 has been shown to associate with Wnt (34), calcineurin-NFAT (35) and protein kinase C (PKC) - transient receptor potential canonical 7 (TRPC7) signaling (36), we also analyzed proteins in these pathways. Immunoblotting showed that the TRPC7 protein level was reduced in both sedentary and exercised syndecan-4^-/-^ (Fig. 5C, KO-SED and KO-ET versus WT-SED). However, no differences were detected in dishevelled (Dvl), β-catenin, frizzled-7, low-density lipoprotein receptor-related protein 6 (LRP6) or Wnt4 in syndecan-4^-/-^ versus WT, indicating no involvement of canonical Wnt/β-catenin or non-canonical Wnt signaling (Suppl. Fig. 2C-G). Similarly, no differences in the PKC or RCAN4.1 (regulator of calcineurin and a target for NFAT) levels were detected (Suppl. Fig. 2H-I).

Taken together, our data indicate that the syndecan-4^-/-^ muscle had increased activation of the Akt/mTOR/S6K1 and Notch-HES-1 pathways and that these pathways were not induced any further by exercise. Except for an also reduced TRPC7 level, few other changes were observed.

## Discussion

In this study we have characterized the TA muscle from syndecan-4^-/-^ mice and analyzed molecules involved in ECM remodeling and muscle growth, both under steady state conditions and in response to exercise. Compared to WT, we found that syndecan-4^-/-^ mice had reduced body weight, reduced muscle weight, muscle fibers with a smaller cross-sectional area, and reduced expression of myogenic regulatory transcription factors. The syndecan-4-/- mice had also increased activation of the Akt/mTOR/S6K1 and Notch-HES-1 pathways, a reduced TRPC7 level, increased syndecan-2 expression, and altered expression of several ECM components. These molecular changes were observed under steady state conditions, and in contrast to WT mice, were mostly not affected further by exercise.

Closer inspection of the syndecan-4^-/-^ muscle showed that the muscle fibers were on average smaller than wild type fibers, which probably accounts for the reduced TA weight. Consistent with previous findings (20), we found that myoD and myogenin were reduced in syndecan-4^-/-^ mice of 12-22 weeks. MyoD is required for MuSC activation, proliferation and differentiation (37). Myogenin expression is upregulated during differentiation and directs differentiating myoblasts to become terminally differentiated (37). Consistent with a role of syndecan-4 in myogenesis, we have previously also demonstrated that the cytoplasmic part of syndecan-4 plays an important role in the early fusion process of myoblasts *in vitro* (38). Our findings of the reduced body weight, muscle weight, muscle fibers and myogenic regulatory transcription factors in syndecan-4^-/-^ are illustrated in Fig. 6A.

**Fig. 6.**
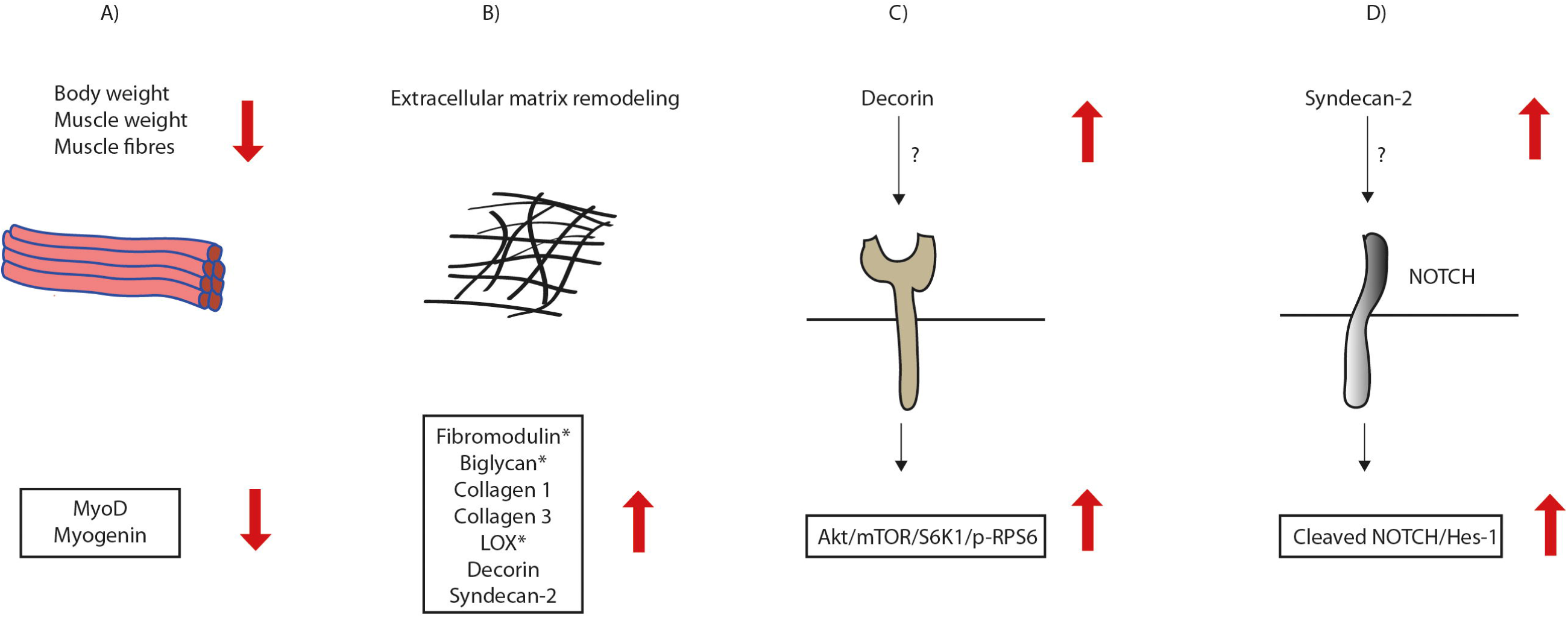
Illustration of alterations in the syndecan-4^-/-^ mice. (**A**) The syndecan-4^-/-^ mice had reduced body weight, muscle weight, muscle fibers and expression of myogenic regulatory transcription factors. (**B**) The syndecan-4^-/-^ muscle had increased mRNA levels of syndecan-2 and the ECM components decorin, fibromodulin, biglycan, collagen 1, collagen 3 and the collagen cross-linking enzyme lysyl oxidase (LOX). However, fibromodulin, biglycan and LOX were reduced at the protein level (denoted with star), suggesting that these ECM components are more susceptible to degradation or less efficiently translated when syndecan-4 is absent. The syndecan-4^-/-^ muscle had also increased activation of the (**C**) Akt/mTOR/S6K1 and (**D**) Cleaved Notch1-HES-1 pathways.

Previous experiments have shown that both syndecan-3 and syndecan-4 are affected by exercise in humans. Syndecan-4 was reported up-regulated after acute exercise, whereas syndecan-3 was up-regulated after long-term exercise (39). However, in another study using rats, exercise did not change the syndecan-4 level (40). In our study using mice subjected to two weeks treadmill running, syndecan-2, but not syndecan-4 was increased, suggesting a potential involvement of syndecan-2 in the muscle response to exercise. Interestingly, we also observed higher levels of syndecan-2 in syndecan-4^-/-^ compared to WT (illustrated in Fig. 6B). Upregulation of syndecan-2 has been reported as a compensatory mechanism during cartilage development in syndecan-4^-/-^ (41). However, syndecan-2 was not higher in the left ventricle from syndecan-4^-/-^ compared to WT (42), suggesting that compensatory upregulation of syndecan-2 is tissue dependent.

Although the mRNA levels of fibromodulin, decorin, biglycan, collagen 1, collagen 3 and the collagen cross-linking enzyme LOX were increased in syndecan-4^-/-^ mice, the protein levels of fibromodulin, biglycan and LOX were reduced (illustrated in Fig. 6B). This finding might suggest that some ECM components are more susceptible to degradation or less efficiently translated when syndecan-4 is absent. Indeed, proteins involved in the translational machinery have been identified in the cardiac syndecan-4 interactome (33, 43), supporting a role for syndecan-4 in translation. Decorin is abundant in adult skeletal muscle and expressed during skeletal muscle differentiation (18), and consistent with our mice mRNA data, reported to be upregulated in rats after moderate treadmill running (44). Biglycan expression is reported highly upregulated during injury-induced regeneration, both in growing myotubes and in activated precursor cells, then the expression decreases again during maturation of muscle fibers (45, 46). The relationship between biglycan and syndecan-4 in skeletal muscle is not known, however a recent publication using cultured vascular endothelial cells suggests biglycan as a regulatory molecule of the ALK5-Smad2/3-TGF-β1 signaling, with a direct negative impact on syndecan-4 expression (47). Further experiments are required to investigate whether the ALK5-Smad2/3-TGF-β1 pathway is also involved in the biglycan-syndecan-4 relationship in skeletal muscle. Fibromodulin is also central in the myogenic program and plays an important role in tissue repair (48). Specifically, fibromodulin triggers myoblast differentiation by preventing myostatin binding to its receptor; the activin receptor type-2B. Fibromodulin, biglycan and decorin are also suggested to regulate collagen fibrillogenesis and crosslinking, and thus ECM integrity and function of skeletal muscle (49). Whether the increased mRNA expression of these ECM components, including collagens and LOX, represent a compensatory mechanism to improve ECM function and trigger myoblast differentiation in the syndecan-4^-/-^ muscle, has to be determined in future work. In contrast to our findings, the syndecan-4^-/-^ heart has rather reduced mRNA levels of collagen 1, collagen 3 and LOX following pressure overload, leading to an impaired cross-linking and collagen assembly (19). Our group has also previously shown that syndecan-4 is essential for cardiac myoblast differentiation (50) and that the syndecan-4^-/-^ hearts do not develop concentric hypertrophy, but rather left ventricle dilatation and dysfunction following pressure overload (35). Although a syndecan-4-calcineurin-NFAT pathway appeared essential for these events in the heart (35, 50, 51), we found no changes in the calcineurin-NFAT pathway in the syndecan-4^-/-^ TA muscle. Thus, although syndecan-4 is involved in both differentiation and cell growth of the heart and skeletal muscle, different pathways seem to be involved.

We also found higher levels of pSer473-Akt and pSer235/236-RPS6 in syndecan-4^-/-^. The Akt/mTOR pathway is an important signaling pathway promoting muscle growth and fibrosis (52, 53). Akt induces protein synthesis and hypertrophy by activating the serine/threonine kinase mTOR and the ribosomal protein S6 kinase 1 (SK61) (29). SK61 is one of the main kinases phosphorylating ribosomal protein 6 (RPS6) (54). Although RPS6 is primarily a structural protein of the ribosome, phosphorylation of RPS6 has been shown to be directly involved in cell size control (31), and affects both translation as well as transcription efficiency. Our finding of an increased pSer235/236-RPS6 level in the syndecan-4^-/-^ mice was quite surprising, since the smaller cell size and smaller cross-sectional area of myofibers rather links to a reduced pSer235/236-RPS6 level (30, 31). Our finding of the increased activation of the Akt/mTOR/S6K1 pathway in syndecan-4^-/-^ is illustrated in Fig. 6C. Notably, decorin, which was also increased in syndecan-4^-/-^, has been shown to activate Akt downstream of IGF1R, in addition to increase myoD and myogenin expression and promote myoblast differentiation (28).

Our findings of higher levels of both Cleaved Notch and HES-1 in syndecan-4^-/-^ mice, strongly suggest an increased Notch signaling when syndecan-4 is absent. HES-1 signaling was also higher in exercised compared to sedentary WT mice. Notch is a regulator of myogenesis and is increased upon physiological stimuli such as exercise (see review (31)). Notch signaling is indispensable for maintaining quiescence and self-renewal of MuSCs (30, 31), and promotes proliferation of MuSCs by keeping low MyoD levels and prevents differentiation (55, 56). Upon ligand binding, the intracellular domain of the Notch receptor is cleaved off (Cleaved Notch) and translocated to the nucleus where it activates the transcription factor CLS/RBPJ and induces expression of the transcriptional repressors of the HES and HEY family (45, 46). Interestingly, both syndecan-2 and 3 have been linked to Notch signaling pathways (43, 57). Syndecan-3 interacts with Notch at the plasma membrane in the MuSC, and this interaction has been suggested to be important for Notch cleavage (57). Loss of syndecan-3 impairs Notch signaling, alters MuSC homeostasis and leads to progressively increased myofiber size, which is especially noticed in repeatedly injured and dystrophic mice (11, 57). Syndecan-2 is reported highly abundant in early-differentiated myoblasts (19). Interestingly, Cleaved Notch has been shown to induce syndecan-2 expression in smooth muscle cells, which again interacts with the Notch receptor at the surface membrane and promotes further Notch signaling (32). Whether the increased Notch signaling we observed in syndecan-4^-/-^ is due to the increased syndecan-2 level or *vice versa* has to be determined by future experiments (illustrated in Fig. 6D).

Desmin was the only protein in our analyses that was increased in exercised WT, but neither changed in sedentary or exercised syndecan-4^-/-^. Desmin is the most abundant intermediate filament protein in adult skeletal muscle, expressed at low levels in MuSCs and increased during myogenic commitment and differentiation (27). Human heterozygous desmin mutations link to muscular weakness and different skeletal and cardiac myopathies. In line with this, the desmin^-/-^ mouse model is also reported weaker; it fatigued more easily and showed an impaired performance on endurance tests (58). Whether the syndecan-4^-/-^ mice also fatigues more easily must be determined in larger endurance tests in future.

## Conclusions

Conclusively, our data show that syndecan-4^-/-^ mice had a lower body weight, lower TA muscles weight, smaller muscle fibers, reduced myoD and myogenin expression, changes in several ECM components, an increased syndecan-2 expression, a reduced TRPC7 level, and increased activation of the Akt/mTOR/S6K1 and Notch-HES-1 pathways. These molecular changes were observed under steady state conditions, and in contrast to WT, were mostly not affected by exercise. Altogether our data suggest an important role of syndecan-4 in muscle development.

## Declarations

### Ethics approval and consent to participate

All animal experiments were performed in accordance with the National Regulation on Animal Experimentation in accordance with an approved protocol and the Norwegian Animal Welfare Act and conform the NIH guidelines

### Consent for publication

Not applicable

### Availability of data and material

All data generated or analysed during this study are included in this published article

### Authors’ contributions

S.B.R: Conceptualization, Investigation, Methodology, Visualization, Writing the original draft; C.R.C: Conceptualization, Investigation, Methodology, Visualization, Writing the original draft; J.M.A Conceptualization, Investigation, Methodology, Review and editing the original draft; A.P: Conceptualization, Visualization, Review and editing the original draft; V.H: Investigation, Methodology, Review and editing the original draft; M.L: Investigation, Methodology, Review and editing the original draft; K.H.L: Investigation, Methodology, Review and editing the original draft; I.S: Methodology, Review and editing the original draft, Funding acquisition S.O.K: Conceptualization, Investigation, Review and editing the original draft. G.K; Conceptualization, Investigation, Review and editing the original draft, Funding acquisition; M.E.P; Conceptualization, Investigation, Methodology, Visualization, Writing the original draft.

## Acknowledgements

We would like to thank Anita Kaupang for some of the immunoblot analyses and Per Kristian Lunde for help with muscle dissection.

## List of abbreviations

MuSC: skeletal muscle stem cells;
ECM: extracellular matrix;
MRFs: myogenic regulatory transcription factors;
MyoD: myoblast determination protein 1;
Myf: myogenic factor;
TA: tibialis anterior;
SED: sedentary;
WT: wild type;
ET: exercised trained;
IGF1R: insulin-like growth factor I receptor;
RPS6: ribosomal protein 6;
SK61: ribosomal protein S6 kinase 1;
PKC: protein kinase C;
GAG: glycosaminoglycan;
SLRPs: small leucine-rich proteoglycans;
TRPC7: Transient Receptor Potential Cation Channel Subfamily C Member 7;
Dvl: dishevelled;
LRP6: Low-density lipoprotein receptor-related protein 6;
LOX: Lysyl oxidase.

