## Supplementary figures and images for "Syndecan-4^-/-^ mice have smaller muscle fibers, increased Akt/mTOR/S6K1 and Notch/HES-1 pathways, and alterations in extracellular matrix components"

### Supplementeray figure 1

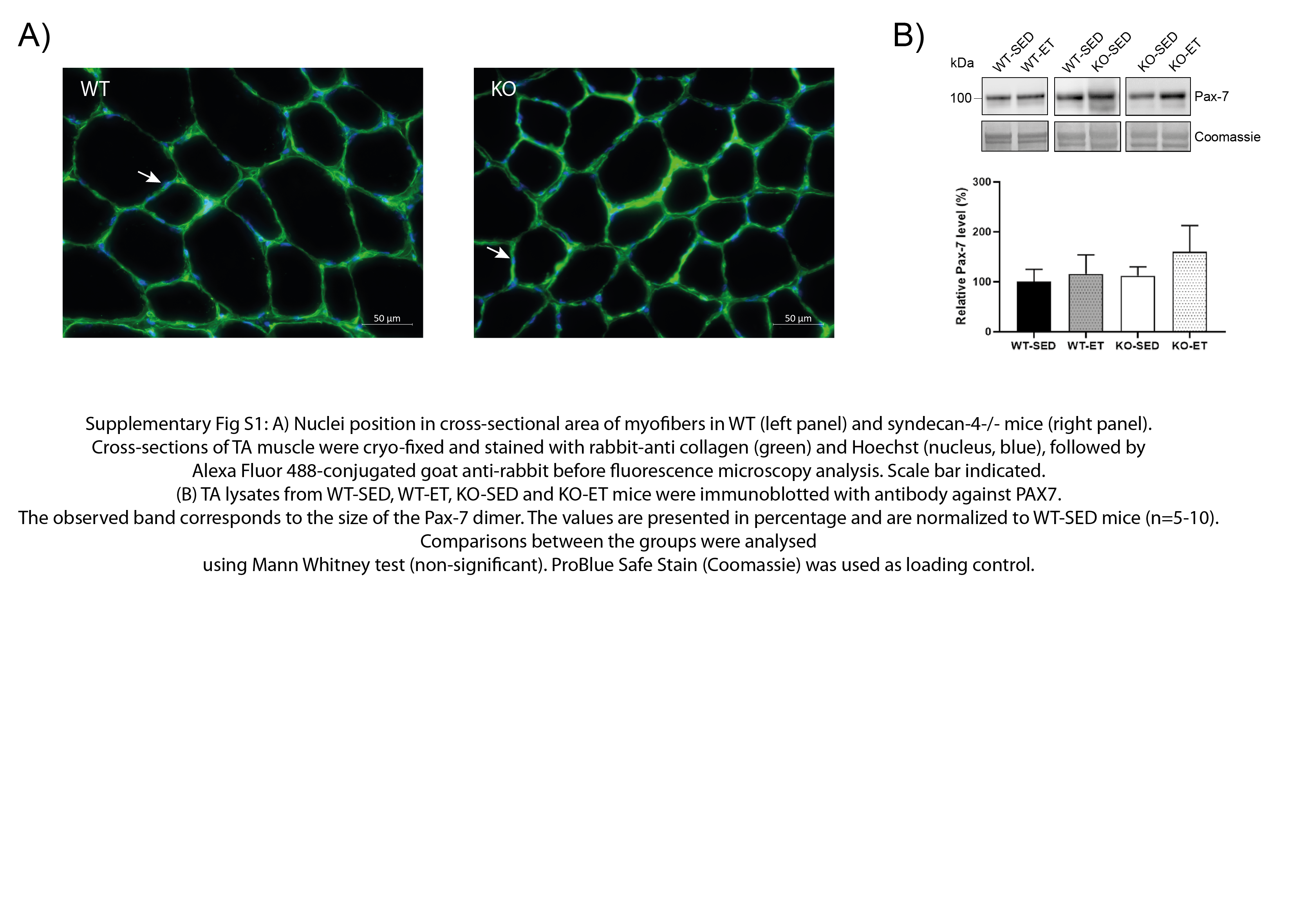

### Supplementeray figure 2

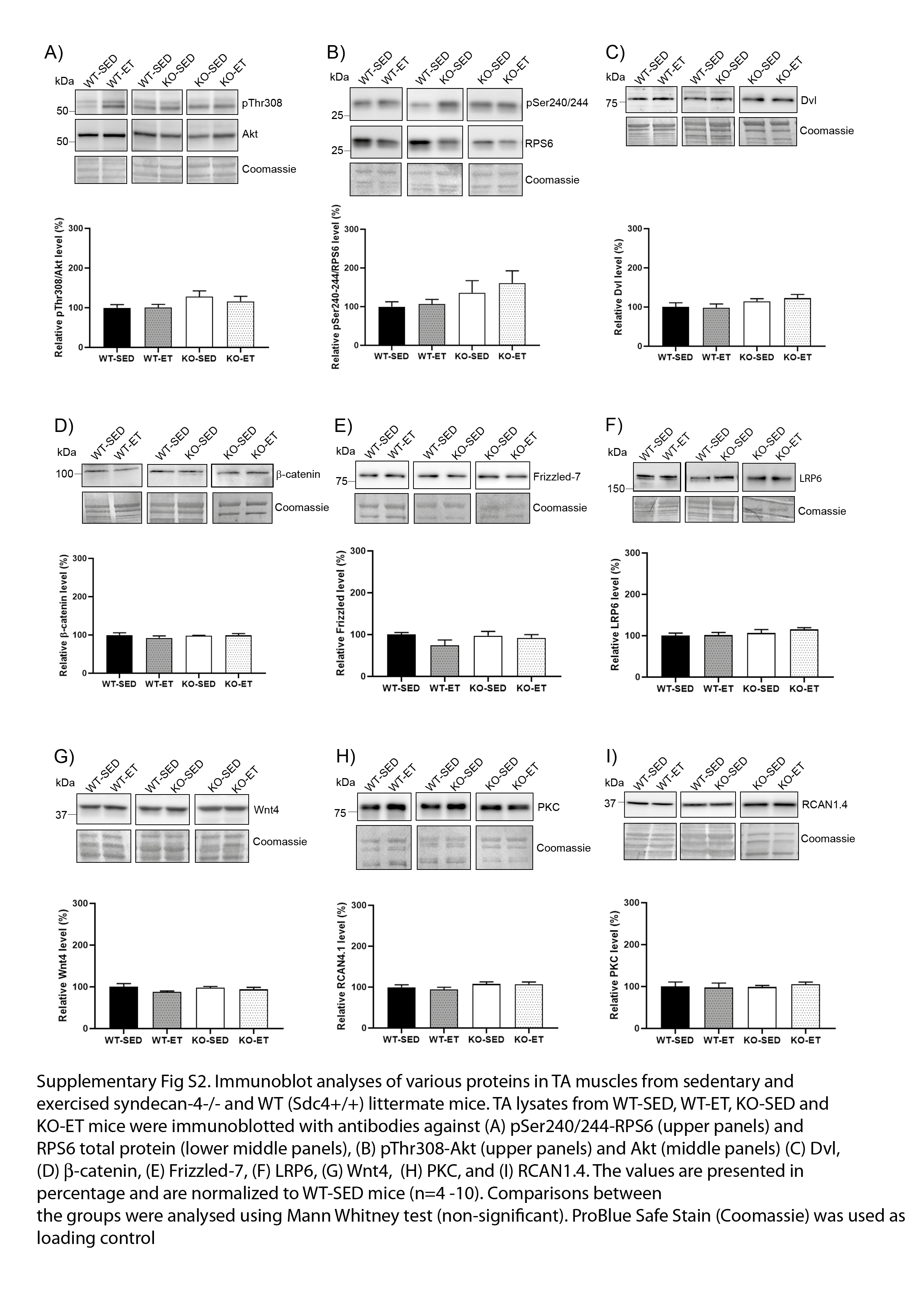
